# Variation in histone configurations correlates with gene expression across nine inbred strains of mice

**DOI:** 10.1101/2022.11.02.514900

**Authors:** Anna L. Tyler, Catrina Spruce, Romy Kursawe, Annat Haber, Robyn L. Ball, Wendy A. Pitman, Alexander D. Fine, Narayanan Raghupathy, Michael Walker, Vivek M. Philip, Christopher L. Baker, J. Matthew Mahoney, Gary A. Churchill, Jennifer J. Trowbridge, Michael L. Stitzel, Kenneth Paigen, Petko M. Petkov, Gregory W. Carter

## Abstract

It is well established that epigenetic features, such as histone modifications and DNA methylation, are associated with variation in gene expression across cell types. Less well known is the extent to which epigenetic states vary across genetically diverse individuals, and whether such variation corresponds to inter-individual variation in gene expression. To investigate genetically driven variation in epigenetics, we conducted a survey of epigenetic modifications and gene expression in hepatocytes of nine inbred mouse strains. We surveyed four histone modifications (H3K4me1, H3K4me3, H3K27me3, and H3K27ac), and DNA methylation. We used ChromHMM to identify 14 chromatin states, each of which represented a distinct combination of the four histone modifications. We found that chromatin states varied widely across the nine strains and that epigenetic state was strongly correlated with local gene expression. We replicated this correspondence between chromatin state and gene expression in an independent population of Diversity Outbred mice in which we imputed local chromatin state. In contrast, we found that DNA methylation did not vary across the inbred strains and was not correlated with variation in gene expression in DO mice. This work suggests that chromatin state is highly influenced by local genotype and may be a primary mode through which expression quantitative trait loci (eQTLs) are mediated. Through examples, we demonstrate that naturally occurring chromatin state variation, in conjunction with gene expression, can aid in functional annotation of the mouse genome. Finally, we provide a data resource that documents variation in chromatin state in hepatocytes across genetically distinct mice.

## Introduction

Although different cell types in an organism house the same genome, each has a distinct pattern of gene expression. These cell-type specific patterns of expression are established through epigenetic modification of both histones (Xu *et al*., 2021; Godini *et al*., 2018) and DNA methylation (Wiench *et al*., 2011; Ji *et al*., 2010), which influence the accessibility of DNA to transcription machinery (Lawrence *et al*., 2016; Jones, 2012; Moore *et al*., 2013).

This variation in epigenetic landscapes across cell types has been extensively documented (Ernst *et al*., 2011; Kundaje *et al*., 2015) and has been used to richly annotate functional elements in both mouse (Stamatoyannopoulos *et al*., 2012; Baker *et al*., 2019; Yue *et al*., 2014) and human genomes (Kundaje *et al*., 2015; Ernst and Kellis, 2013, 2010). Such functional annotations provide insight into mechanisms of gene regulation. Because the majority of disease-associated genetic variants discovered in humans are in gene regulatory regions, it has been suggested that it is the regulation of gene expression, rather than alteration of protein function, that is the primary mechanism through which genetic variation confers disease risk (Maurano *et al*., 2012; Farh *et al*., 2015; Pennisi, 2011; Hindorff *et al*., 2009). Detailed epigenomic landscapes, therefore, may provide important mechanistic insight linking genotype to disease risk.

However, although variation in epigenetic modifications across cell types has been deeply explored, we have relatively little knowledge of inter-individual variation in epigenetic modifications. Does genetic variation across individuals influence the epigenetic landscape? To what extent does this variation result in altered gene expression? The generation of a more complete picture of interindividual variation in epigenetic modifications will increase our understanding of the mechanisms of gene regulation, provide insights into the mechanisms establishing cell type-specific epigenetic landscapes, and improve the functional annotation of the genome as it relates to the regulation of gene expression and disease risk.

To investigate the effect of genetic variation on epigenetic variation, we performed a survey of epigenetic variation in hepatocytes across nine inbred mouse strains. We included the eight founders of the Diversity Outbred/Collaborative Cross (DO/CC) (Svenson *et al*., 2012) mice, as well as DBA/2J, which, along with C57BL/6J, is one of the founders of the widely used BxD recombinant inbred panel of mice (Ashbrook *et al*., 2019). We assayed four histone modifications: H3K4me3, which is associated with promoter regions (Heintzman *et al*., 2007; Bernstein *et al*., 2005), H3K4me1, which is associated with enhancer regions (Heintzman *et al*., 2007), H3K27me3, which is associated with polycomb repression (Bonasio *et al*., 2010), and H3K27ac, which has been associated with enhancement of active enhancers and promoters (Creyghton *et al*., 2010; Heintzman *et al*., 2009; Rada-Iglesias *et al*., 2011). We also assayed DNA methylation which is associated with repression of expression (Moore *et al*., 2013; Jones, 2012).

We used ChromHMM (Ernst and Kellis, 2012) to identify 14 chromatin states, each representing a unique combination of the four histone marks. We investigated the association between variation in these states and variation in gene expression across the nine strains. We separately investigated the relationship between DNA methylation and gene expression across strains.

We further investigated the relationship between epigenetic state and gene expression by imputing the 14 chromatin states and DNA methylation into a population of DO mice. We then mapped gene expression to the imputed epigenetic states to assess the extent to which eQTLs were driven by variation in epigenetic modification.

## Results

Both gene expression and epigenetic state were consistent within mouse strains but varied across the strains suggesting strong genetic regulation of both modalities. This is seen as a clustering of individuals from the same strain in principal component plots of transcriptomic and epigenetic features (Figure 1). Patterns of gene expression (Figure 1A), DNA methylation (Figure 1B) and individual histone modifications (Figure 1C-F) clustered in similar patterns although a relatively small percent of the variation in the methylome was related to strain. The three subspecies *musculus* (in red), *castaneous* (in green) and *domesticus* (all others) were widely separated suggesting that subspecies structure made up the majority of the observed variance. The domesticus strains largely clustered together. These data provide evidence that epigenetic features relate to gene expression in a manner that is consistent with the subspecific origin of the mouse strains.

**Figure 1:**
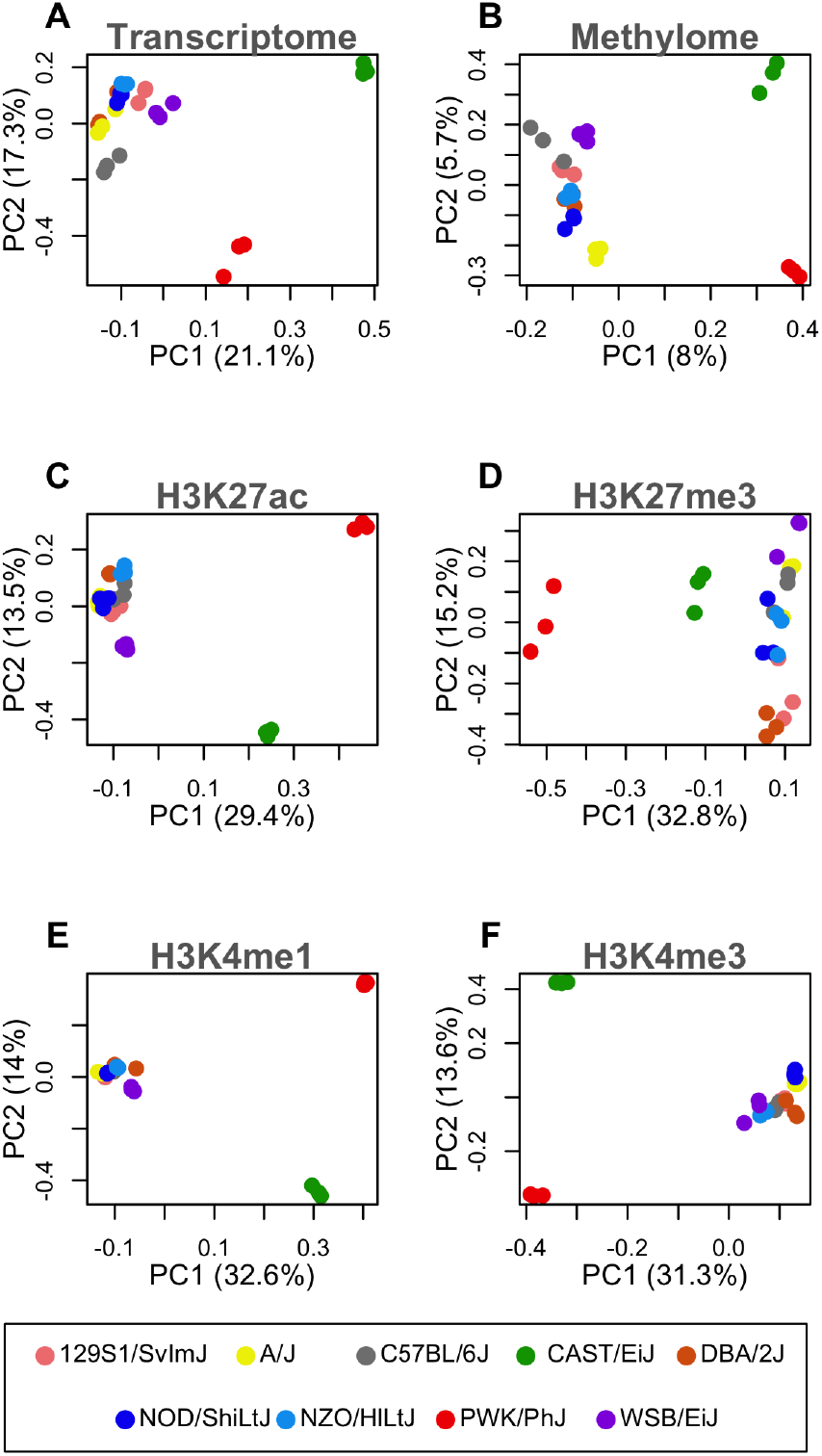
The first two principle components of each genomic feature across nine inbred strains of mouse. In all panels each point represents an individual mouse, and strain is indicated by color as shown in the legend at the bottom of the figure. Each panel is labeled with the data used to generate the PC plot. (A) Hepatocyte transcriptome - all transcripts sequenced in isolated hepatocytes. (B) DNA methylation - the percent methylation at all CpG sites shared across all individuals. (C-F) Histone modifications - the peak heights of the indicated histone modification for sites shared across all individuals.

### Chromatin state overview

To investigate the association between histone modifications and gene expression, we used ChromHMM to identify 14 chromatin states composed of unique combinations of the four histone modifications. Panel A in Figure 2 shows the representation of each histone modification across the states.

**Figure 2:**
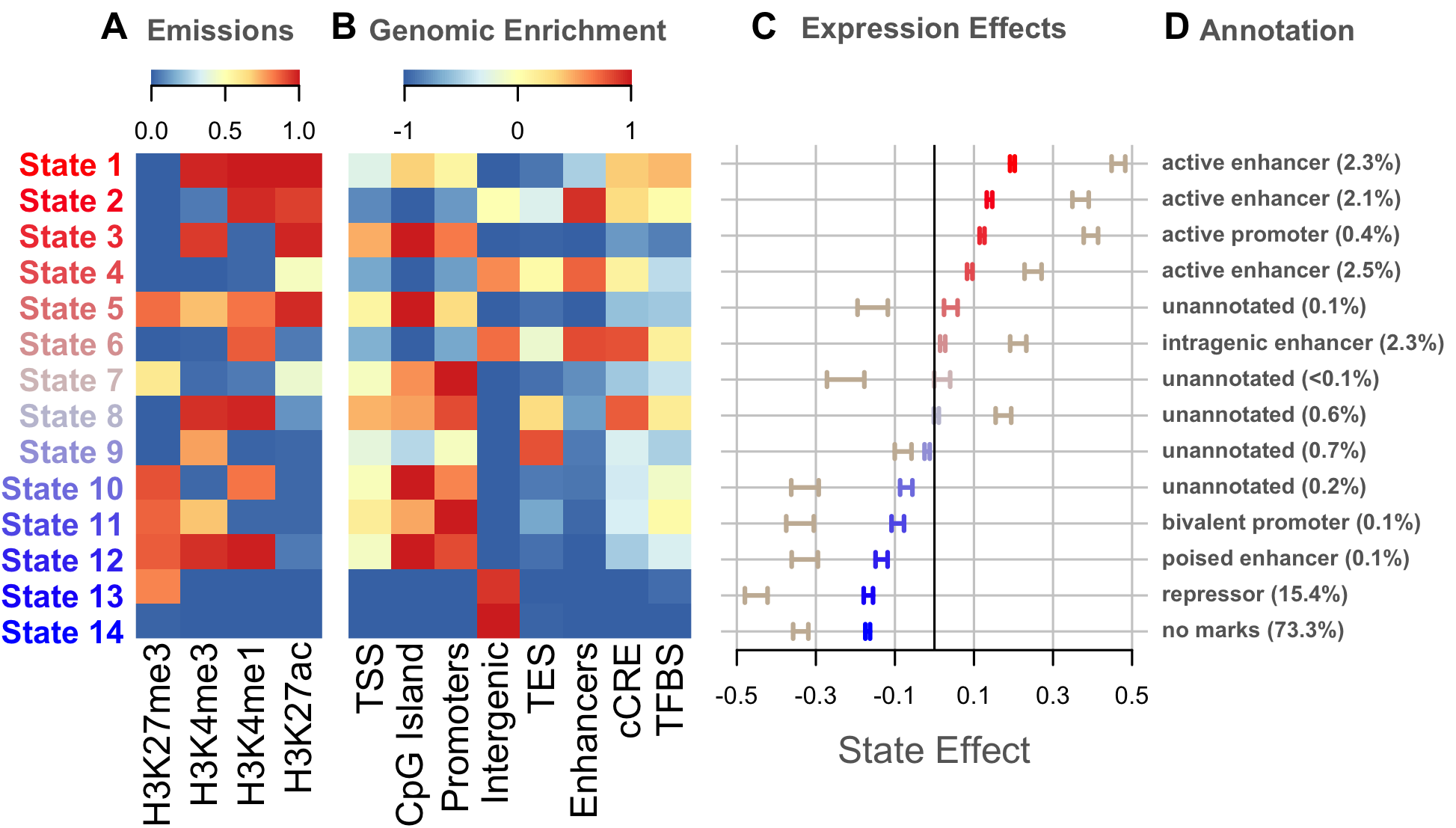
Overview of chromatin state composition, genomic distribution, and effect on expression. (A) Emission probabilites for each histone modification in each chromatin state. Blue indicates the absence of the histone modification, and red indicates the presence of the modification. (B) The distribution of each state around functional elements in the genome. Red indicates that the state is enriched near the annotated functional element. Blue indicates that the state is depleted near the annotated functional element. Abbreviations are as follows: TFBS = transcription factor binding sites, cCRE = candidate cis-regulatory element, TSS = transcription start site, TES = transcription end site. (C) The effect of variation in the state on gene expression. Bars are colored based on the size and direction the state’s effect on expression. Darker bars show the effects on expression of chromatin state variation across strains. Tan bars show the effects on expression of chromatin state variation across genes. (D) Plausible annotations for each state based on combining the data in the previous three panels. The numbers in parentheses indicate the percent of the genome that was assigned to each state.

The states were enriched around known functional elements in the mouse genome (Figure 2B). For example, the majority of the states were enriched around transcription start sites (TSS), and other TSS-related functional elements, such as promoters and CpG islands. Two states (States 13 and 14) were primarily found in intergenic regions. Three states (States 2, 6, and 4) were enriched around known enhancers, and one (State 9) was enriched predominantly near the transcription end sites (TES).

Most states were also associated with variation in gene expression across strains (Figure 2C). The states in Figure 2 are shown in order of their effect on expression, which helps illustrate several patterns. The state with the largest negative effect on gene expression, State 14, is the absence of all measured modifications. Other states associated with reduced gene expression contained the repressive mark H3K27me3. The states with the largest positive effects on expression all had some combination of the activating marks, H3K4me3, H3K4me1, and H3K27ac. The repressive mark was less commonly seen in these activating states. These global patterns of positional enrichment and association with expression largely agree with previous findings.

By merging the information from Figure 2A-C, we were able to suggest annotations for many of the 14 chromatin states (Figure 2D). States with the strongest effects on expression had the clearest annotations, while states with weaker effects remained unannotated.

### Spatial distribution of epigenetic modifications around gene bodies

In addition to looking for enrichment of chromatin states near annotated functional elements, we characterized the fine-grained spatial distribution of each state around gene bodies by normalizing gene position to run from 0 at the TSS to 1 at the TES (See Methods) (Figure 3A-B). We similarly characterized the distribution of CpG sites and their percent methylation at this gene-level scale (Figure 3C-D).

**Figure 3:**
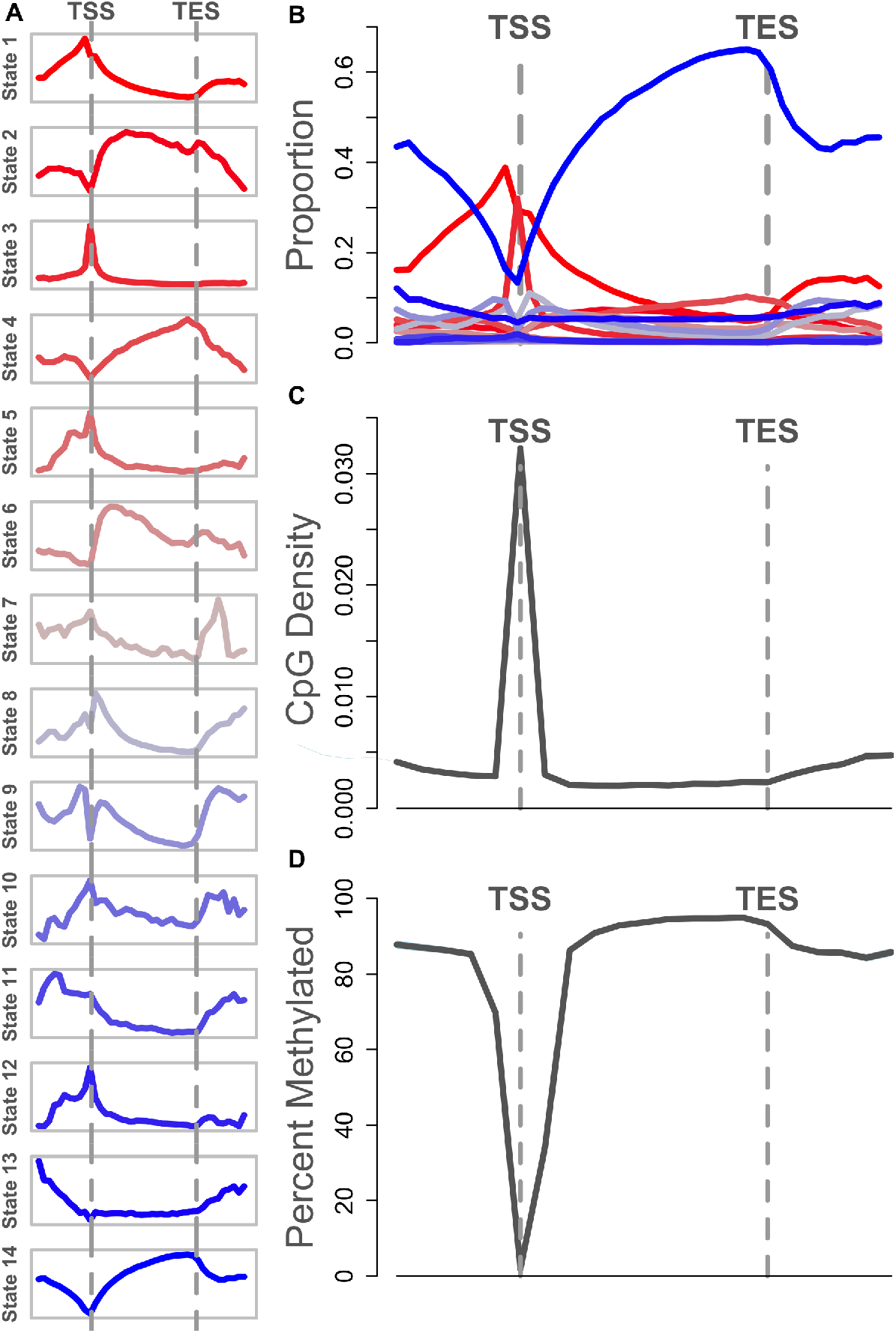
Relative abundance of chromatin states and methylated DNA. (A) Each panel shows the abundance of a single chromatin state relative to gene TSS and TES. The *y*-axis in each panel is the proportion of genes containing the state. Each panel has an independent *y*-axis to better show the shape of each curve. The *x*-axis is the relative gene position. The TSS and TES are marked as vertical gray dashed lines. (B) The same data shown in panel A, but with all states overlayed onto a single set of axes to show the relative abundance of the states. (C) The density of CpG sites relative to the gene body. The *y*-axis shows the number of CpG sites per base pair. The density is highest near the TSS. CpG sites are less dense within the gene body and in the intergenic space. (D) Percent methylation relative to the gene body. The *y*-axis shows the median percent methylation at CpG sites, and the *x*-axis shows relative gene position. CpG sites near the TSS are unmethylated relative to intragenic and intergenic CpG sites.

The spatial patterns of the individual chromatin states are shown in (Figure 3A), and an overlay of all states together (Figure 3B) emphasizes the difference in abundance between the most abundant states (States 1, 3, and 14), and the remaining states, which were relatively rare.

Each chromatin state had a characteristic distribution pattern. For example, State 14, which was characterized by the absence of all measured histone modifications, was strongly depleted near the TSS, indicating that this region is commonly subject to the histone modifications we measured here. In contrast, States 1 and 3 were both enriched at the TSS. State 3 was very narrowly concentrated right at the TSS, whereas State 1 was more broadly abundant both upstream and downstream of the TSS. Both were associated overall with increased expression across inbred mice (indicated by red shading in Figure 3), suggesting promoter or enhancer functions. The third state in this group of expression-enhancing states, State 2, was depleted nere the TSS, but enriched within the gene body, suggesting that this state may mark active intragenic enhancers.

States with weaker effects on expression (indicated by grayer shades in Figure 3) were of lower abundance, but had distinct distribution patterns around the gene body suggesting the possibility of distinct functional roles in the regulation of gene expression.

DNA methylation showed similarly characteristic variation in abundance (Figure 3C-D). The TSS had densely packed CpG sites relative to the gene body (Figure 3C). As expected, the median CpG site near the TSS was consistently hypomethylated relative to the median CpG site in intra- and intergenic regions (Figure 3D). All genes used in this analysis were expressed and thus had some degree of hypomethylation.

### Spatially resolved effects on gene expression

The distinct spatial distributions of the chromatin states and methylated CpG sites around the gene body raised the question as to whether the effects of these states on gene expression could also be spatially resolved. To investigate this possibility we tested the association between gene expression and both chromatin state and DNA methylation using spatially resolved models (Methods). We tested the effect of each chromatin state on expression across genes within hepatocytes (Figure 4A) and the effect of each chromatin state on the variation in gene expression across strains (Figure 4B). All chromatin states demonstrated spatially dependent effects on gene expression within hepatocytes. For many of the states, the effects on expression were concentrated at or near the TSS, while in the other states effects were seen across the whole gene. The direction of the effects matched the overall effects of each state seen previously (Figure 2). The spatial effects were recapitulated for almost every state when we measured across strains. That is, chromatin states that either enhanced or suppressed gene expression across hepatocyte genes were similarly related to variation in expression across strains. This suggests that the genetic differences between strains modify chromatin activity in a manner similar to that used across genes. One notable exception was State 6, whose presence upregulated genes within hepatocytes, but did not contribute to expression variation across strains. We also examined the effect of percent DNA methylation across genes within hepatocytes and across strains (Figure 5). As expected, methylation at the TSS was associated with lower expression in hepatocytes. However, percent DNA methylation did not contribute at all to expression variation across strains. This was in part due to an overall lack of variation in percent DNA methylation at the TSS. These results imply that percent DNA methylation does not vary significantly between strains, at least in hepatocytes, and does not contribute to variation in gene expression across genetically diverse individuals.

**Figure 4:**
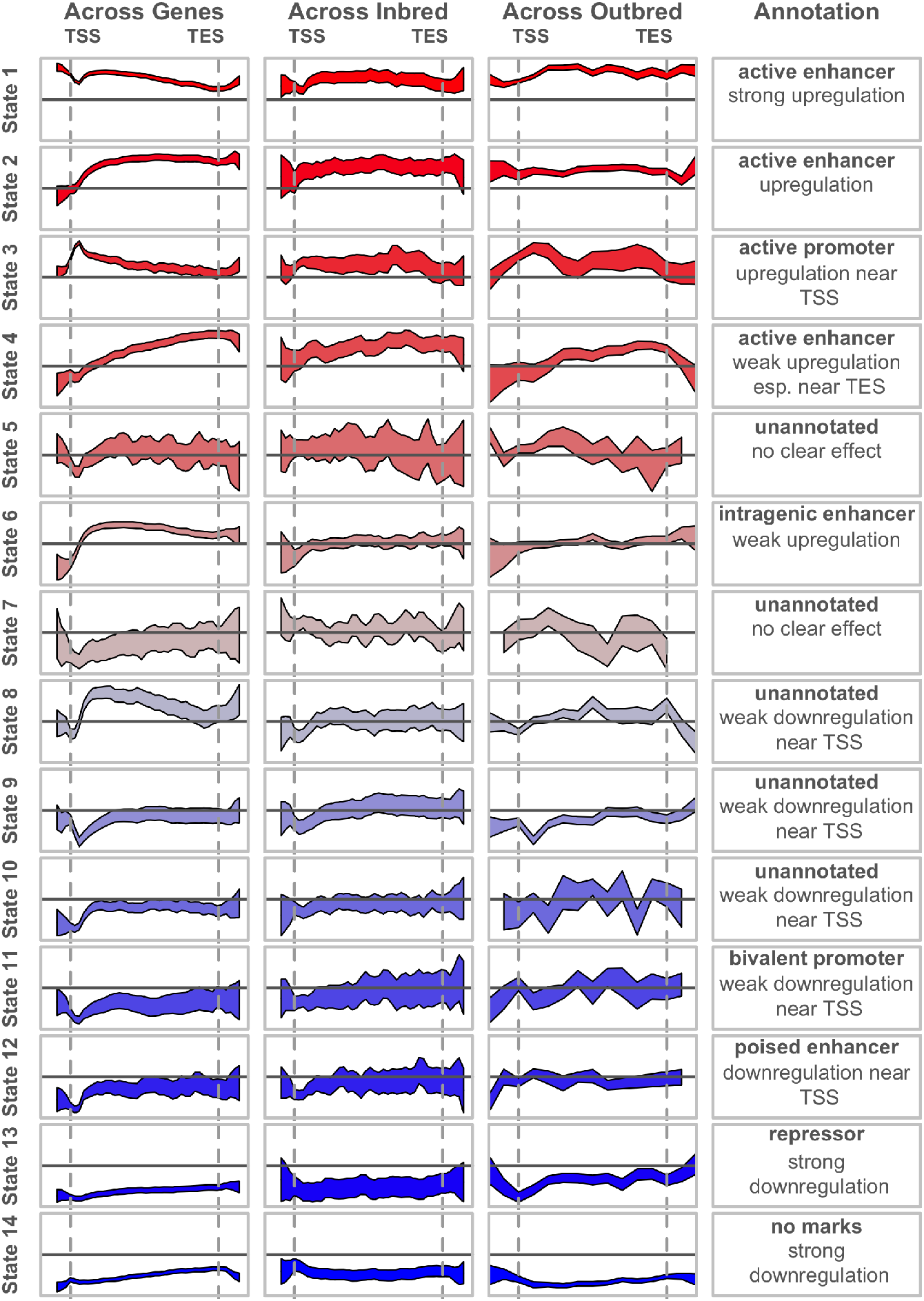
Effects of chromatin states on gene expression. Each column shows the effect of the chromatin states on gene expression in a different experimental context. The first column shows that the presence of some states is correlated with higher abundance genes while the presence of other states is correlated with lower abundance genes in mouse hepatocytes. The second column shows that variation in chromatin state across strains is correlated with variation in gene expression. The third column shows the effect of imputed chromatin state on gene expression in a population of DO mice. The *y*-axis in each column is constant for comparison of states to each other within a single context. The *y*-axes vary across columns to highlight the similarity of the shape of each curve across settings. The final column shows the annotation of each state. All *y*-axes show the *β* coefficient from the linear model shown in the equation. All *x*-axes show the relative position along the gene body. Vertical gray dashed lines mark the TSS and TES.

**Figure 5:**
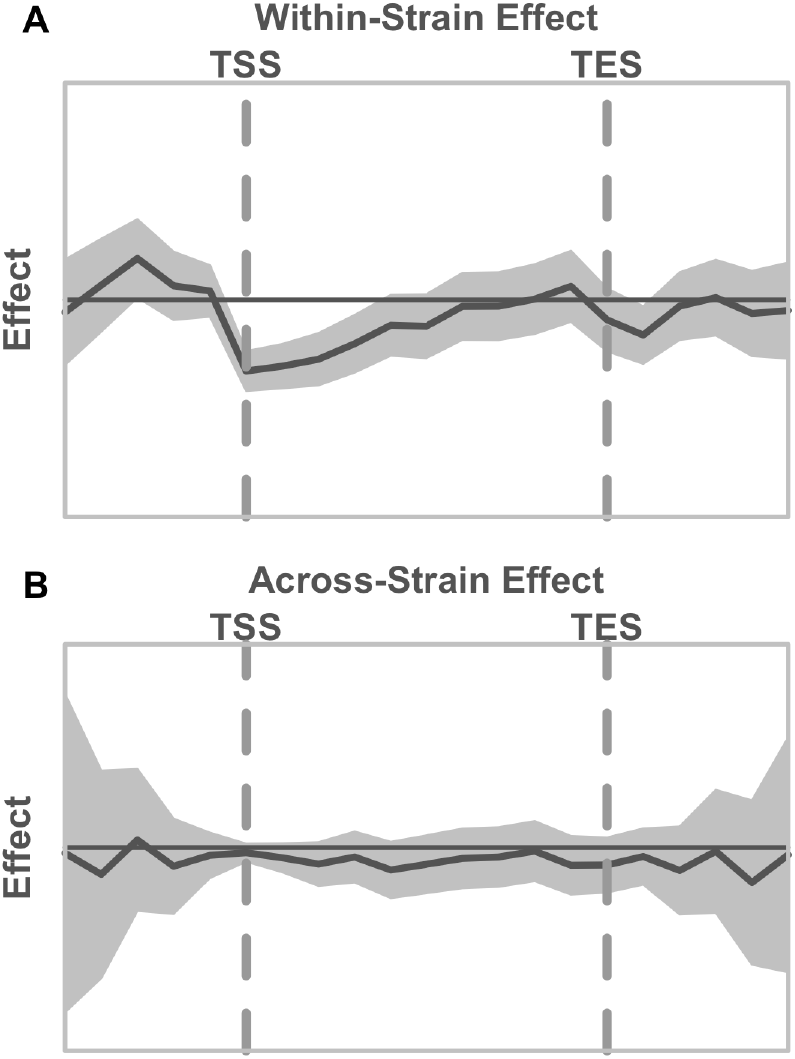
Effect of DNA methylation on gene expression (A) across gene expression in hepatocytes and (B) across inbred strains. The dark gray line shows the estimate of the effect of percent DNA methylation on gene expression. The *x*-axis is normalized position along the gene body running from the transcription start site (TSS) to the transcription end site (TES), marked with vertical gray dashed lines. The horizontal solid black line indicates an effect of 0. The shaded gray area shows 95% confidence interval arond the model fit.

### Imputed chromatin state explained expression variation in diversity outbred mice

Thus far, we have used inbred strains of mice to identify correlations between local chromatin state and gene expression. To compare the contribution of genetic and epigenetic features to expression quantitative trait loci (eQTLs) in a gentically diverse population, we imputed chromatin state, DNA methylation, and SNPs into DO mice (Methods). Chromatin state is largely determined by local genotype, especially early in life (Fraga *et al*., 2005), and can thus be reliably imputed from local genotype. Further, we have shown here that local chromatin state correlates with variation in gene expression across inbred strains. DNA methylation, on the other hand, is known not to be highly heritable (Villicaña and Bell, 2021), and thus cannot be reliably imputed from local genotype. We have also shown here that DNA methylation is not correlated with variation in gene expression across inbred strains. The imputation of DNA methylation thus serves as a negative control–an estimate of a lower bound the ability of a feature imputed from local haplotype to explain gene expression in a new population.

After imputing each genomic feature into the DO population, we mapped gene expression to the imputed features and calculated the variance explained (Methods). The overall distributions of variance explained by each feature across all transcripts are shown in Figure 6.

**Figure 6:**
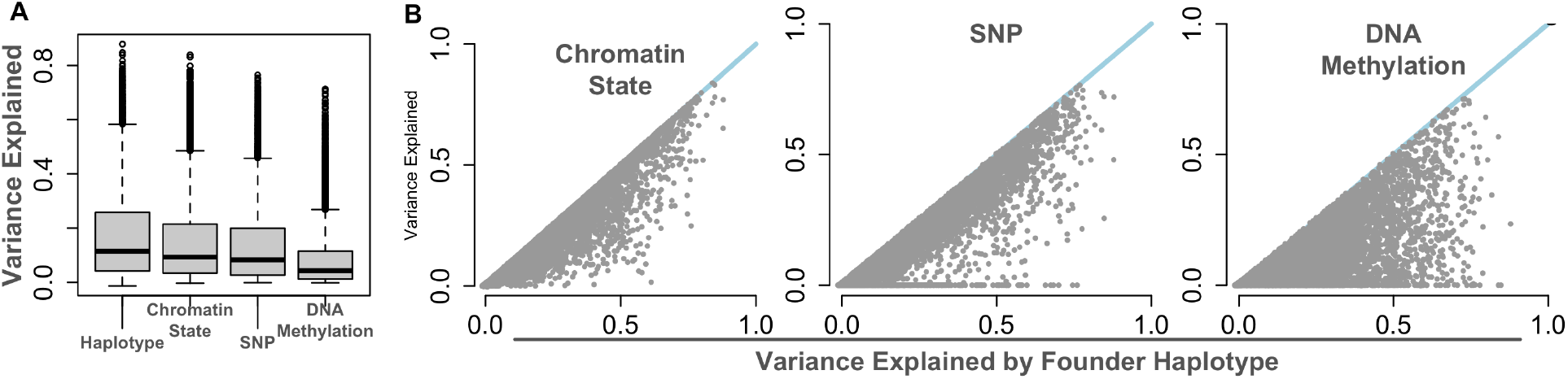
Gene expression variance in a DO population explained by chromatin state compared to three other genomic features: local haplotype, local SNP genotype, and local imputed DNA methylation status. A. Distributions of gene expression variance explained by each feature. B. Direct comparisons of variance explained by local haplotype, and each of the other genomic features. Blue lines show *y* = *x*. Each point is a single transcript.

These distributions show the haplotype effect for the marker nearest each transcript compared with the maximum effect across the gene body for each of the other imputed features. Overall, local haplotype explained the largest amount of variance of gene expression in the DO (*R*^2^ = 0.17; 99% CI = 0.166-0.174). The variance explained by local chromatin state was very highly correlated with that of haplotype (Pearson *r* = 0.96, 99% CI = 0.958-0.962) and explained almost as much variance in gene expression in the DO as local haplotype (*R*^2^ = 0.15, 99% CI = 0.143-0.151). The mean variance explained by SNPs was lower (*R*^2^ = 0.13; 99% CI = 0.131-0.138) than that explained by haplotype and was not as highly correlated with local haplotype as chromatin state was (Pearson *r* = 0.93; 99% CI = 0.927-0.933). DNA methylation, the lower bound for variance explained by a feature imputed from local haplotype, explained the lowest amount of expression variance in the DO population (*R*^2^ = 0.09; 99% CI = 0.082-0.088), and had a much lower correlation to haplotype than either chromatin state or SNPs (Pearson *r* = 0.74; 99% CI = 0.728-0.751). Taken together, these results suggest that the majority of haplotype-associated variance is encoded by the chromatin state of the allele. Moreover, this encoding is primarily local and present in the relevant inbred founder strain.

To investigate whether natural variation in chromatin state can be used to functionally annotate the mouse genome, we looked at the relationship between chromatin state and gene expression in individual genes. An example gene, *Pkd2*, is shown in Figure 7. This figure shows chromatin state (Figure 7B), SNP genotype (Figure 7C), and DNA methylation (Figure 7D) status along the gene body of *Pkd2*. It also shows the association of each of these features with gene expression calculated across the whole gene body (Figure 7A).

**Figure 7:**
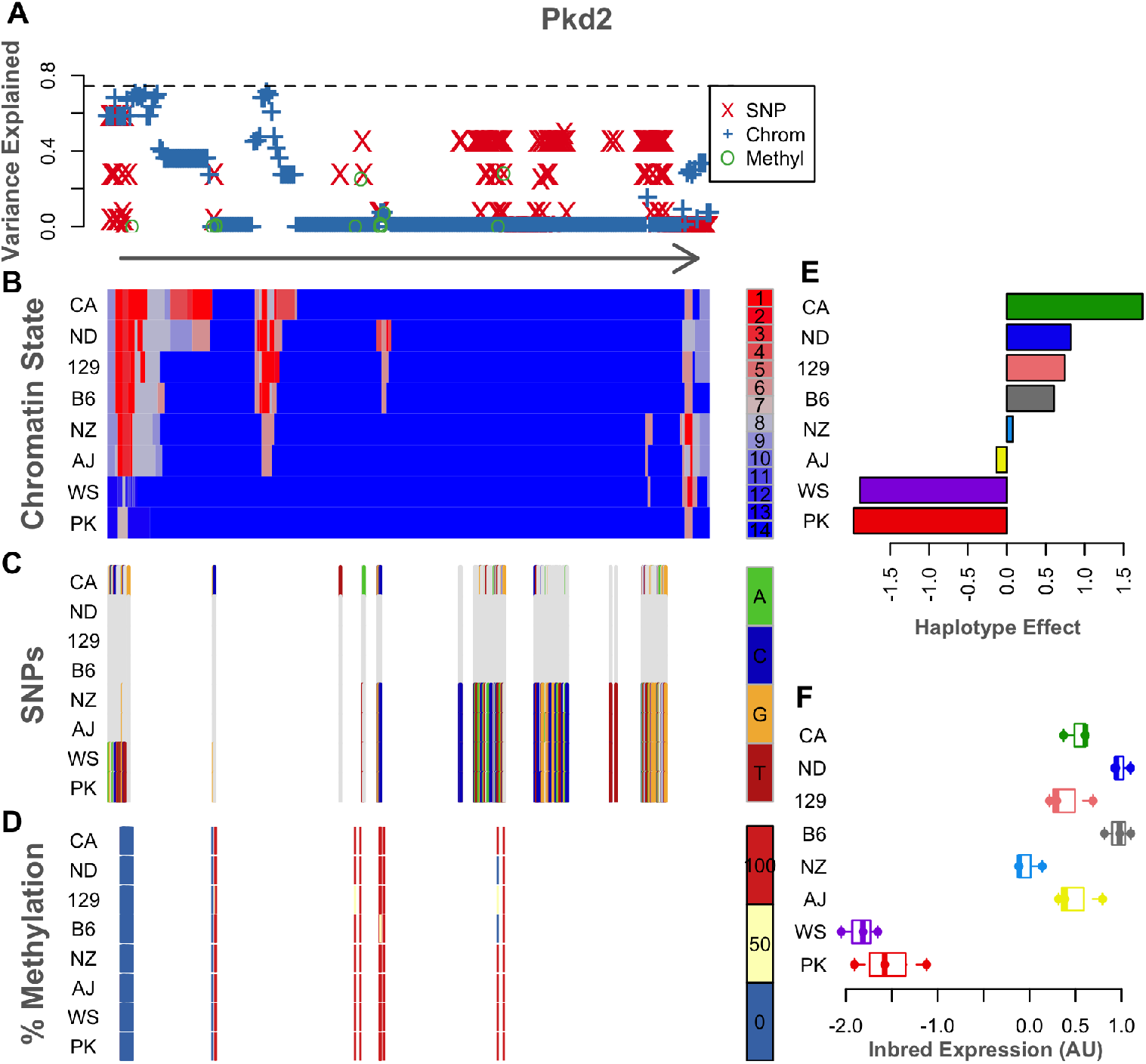
Example of epigenetic states and imuptation results for a single gene, *Pkd2*. The legend for each panel is displayed to its right. (A) The variance in DO gene expression explained at each position along the gene body by each of the imputed genomic features: SNPs - red X’s, Chromatin State - blue plus signs, and Percent Methylation - green circles. The horizontal dashed line shows the variance explained by the haplotype. For reference, the arrow below this panel runs from the TSS of *Pkd2* to the TES and shows the direction of transcription. (B) The chromatin states assigned to each 200 bp window in this gene for each inbred mouse strain. States are colored by their effect on gene expression in the inbred mice. Red indicates a positive effect on gene expression, and blue indicates a negative effect. Each row shows the chromatin states for a single inbred strain, which is indicated by the label on the left. (C) SNPs along the gene body for each inbred strain. The reference genotype is shown in gray. SNPs are colored by genotype as shown in the legend. (D) Percent DNA methylation for each inbred strain along the *Pkd2* gene body. Percentages are binned into 0% (blue) 50% (yellow) and 100% (red). (E) Haplotype effects for expression of *Pkd2* in the DO. Haplotype effects are colored by from which each allele was derived. (F) *Pkd2* expression levels across inbred mouse strains. For ease of comparison, all panels B through F are shown in the same order as the haplotype effects.

The detailed view of this gene identifies two particularly interesting regions. One is at the TSS and the immediately surrounding area, and the other is just downstream of the TSS.

These two regions were both strongly associated expression levels (Figure 7A). Those strains with activating histone states in these regions had much higher expression of *Pkd2* than the two strains that did not have activating states in this region. We hypothesize that these two regions are enhancers.

The spatial patterns in the SNPs only partially mirror those in chromatin state (Figure Figure 7C). SNPs underlying the putative enhancer region at the TSS could potentially influence gene expression by altering local chromatin state. However, the more downstream enhancer has no underlying SNPs, suggesting that there is an alternative mechanism for determining chromatin state at this location. Perhaps SNPs in the TSS region regulate both enhancers.

Percent DNA methylation does not vary across the strains in either of these putative enhancer regions, and does not contribute to variation in expression across genetically distinct individuals (Figure 7D).

## Discussion

In this study we showed that genetic variation across inbred mice alters histone modification patterns in hepatocytes. We further showed that the variation in histone patterns was highly related to variation in gene expression across strains. These observations suggest that within a cell type specific patterns of histone modifications are determined by local genotype, and are likely a major mechanism through which eQTL are generated. This hypothesis was supported by the high concordance between chromatin state, which was imputed from local genotype, and gene expression in an independent outbred population of mice. Thus, across cell types, there is likely a strong interaction between factors determining cell fate, such as transcription factor expression, and local genetics.

In contrast to chromatin state, percent DNA methylation was not associated with variation in gene expression across inbred strains or in the outbred population. At the TSS, this was largely due to a lack of variation in methylation across strains. An example of this observation is shown in panel D of Figure 7. Despite strain variation in both genotype and chromatin state at the TSS of *Pkd2*, DNA methylation was invariant – the CpG island at the TSS is unmethylated in all strains. Thus, although chromatin state appears to be highly influenced by local haplotype, percent DNA methylation is not. CpG islands are highly conserved across vertebrates (Papin *et al*., 2021) and may thus be regions of low genetic variation within this study relative to the surrounding regions in which SNPs and variations in histone modifications are abundant.

Variation in DNA methylation has shown a similar lack of association with gene expression in humans (Villicaña and Bell, 2021). Multiple twin studies have estimated the average heritability of individual CpG sites to be roughly 0.19 (van Dongen *et al*., 2016; Grundberg *et al*., 2013; Bell *et al*., 2012), with only about 10% of CpG sites having a heritability greater than 0.5 (Grundberg *et al*., 2013; Bell *et al*., 2012; McRae *et al*., 2014). Trimodal CpG sites, i.e. those with methylation percent varying among 0, 50, and 100%, have been shown in human brain tissue to be more heritable than unimodal, or bimodal sites (*h*^2^ = 0.8 ± 0.18), and roughly half were associated with local eQTL (Gibbs *et al*., 2010). Here, we did not see an association between trimodal CpG sites and gene expression across strains (Supplemental Figure S5).

The diversity in the effects observed in the 14 chromatin states highlights the importance of analyzing combinatorial states as opposed to individual histone modifications. To illustrate this point, consider the three states with the largest positive effects on transcription. Each of these three states had a distinct combination of the three activatin histone marks: H3K4me1, H3K4me3, and H3K27ac. And although all three states were associated with increased gene expression, each had a distinct spatial distribution. This variation in spatial distribution was mirrored in the spatial effects on transcription. These distinct patterns would not be detectable without analysis of the histone modifications in combination. These results highlight the complexity of the histone code and the importance at analyzing combinatorial states.

While we were able to annotate several states, particularly those with the strongest effects on gene expression, other states were more difficult to annotate. This raises the intruiguing possibility of identifying new modes of expression regulation through histone modification. One of these unannotated states, State 9, had a weak, but consistent negative effect on gene transcription centered within the gene body downstream of the TSS. This state was characterized by high levels of H3K4me3 and low levels of the other three modifications.

The modification H3K4me3 is most frequently associated with increased transcriptional activity (Bernstein *et al*., 2005; Schneider *et al*., 2004; Santos-Rosa *et al*., 2002; Wysocka *et al*., 2006), so the association with state 9 with reduced transcription is a deviation from the dominant paradigm. The physical distribution of this state is also interesting. It was depleted at the TSS, but enriched just upstream and just downstream of the TSS. It was also enriched just downstream of the TES, although it did not appear to influence transcription at this location. The group of genes marked by State 9 were enriched for functions such as stress response, DNA damage repair, and ncRNA processing suggesting that this state may be used to regulate subsets of genes involved in responses to environmental stimuli.

We detected two bivalent states in this survey. Bivalent states are characterized by a combination of activating and repressing histone modifications, and are usually associated with undifferentiated cells (Voigt *et al*., 2013; Vastenhouw and Schier, 2012). Here we identified two bivalent states in adult mouse hepatocytes, and annotated them as a poised enhancer (State 12) and a bivalent promoter (State 11). Both states were associated with downregulation across inbred strains when present near the TSS; however this effect was not replicated in the outbred mice. The lack of replication may be because the effect was too weak to detect given the number of animals in the population.

Both bivalent promoters and poised enhancers are dynamic states that change over the course of differentiation and in response to external stimuli. Bivalent promoters have been studied primarily in the context of development. They are abundant in undifferentiated cells, and are typically resolved either to active promoters or to silenced promoters as the cells differentiate into their final state (Voigt *et al*., 2013; Vastenhouw and Schier, 2012). These promoters have also been shown to be important in the response to changes in the environment. Their abundance increases in breast cancer cells in response to hypoxia (Prickaerts *et al*., 2016). Poised enhancers are also observed during differentiation and in differentiated cells (Bae and Lesch, 2020). In concordance with these previous observations, the genes marked by States 11 and 12 were enriched for vascular development and morphogenesis. That we identified these states in differentiated hepatocytes may indicate that a subset of developmental genes retain the ability to be activated under certain circumstances, such as during liver regeneration in response to damage. It is also possible these states were induced in the inbred strains in respose to stress, rather than genetically coded. This could also explain why the negative effect on gene expression was not replicated in the outbred mice. However, given that we detected this state in all nine inbred strains in relatively equal proportions, this latter hypothesis seems less likely.

The variation of chromatin state across strains, and its correspondence with expression variation offers a unique way to identify gene regulatory regions. Genetic variation serves as a natural perturbation to the regulatory regions, and resulting differences in regulatory annotations can then be linked to variation in gene expression. The *Pkd2* example illustrates how genetic and epigenetic variation can be combined to identify two putative enhancer regions for the gene. The variation in chromatin state further suggests a putative mechanism for the observation of a cis-eQTL at the level of the haplotype.

The discordance between the patterns of chromatin state and SNPs in this gene is particularly interesting. Variation in chromatin state at the intragenic enhancer is present in the absence of local SNPs. This suggests that the presence of the downstream enhancer is determined by another mechanism, perhaps SNPs acting in *trans* to this region, or local variation, such as indels, that was not measured by SNP genotyping. Genetic variation located at a distance from the putative enhancer sites could also potentially alter the 3D configuration of the genome resulting in variable access of transcription factors to the enhancer.

Broadly, local variation in chromatin state was highly correlated with variation in gene expression across individuals, an observation that was replicated in an independent population of genetically diverse, outbred mice. The percent variance explained by chromatin state closely matched that of haplotype, and exceeded that of individual SNPs. These results suggest two things: First, a large portion of the effect of local haplotype on gene expression in mice may be mediated through variation in chromatin state. Second, the intermediate resolution of chromatin state between that of individual SNPs and broad haplotypes carries important imfornation that cannot be resolved at the other levels. Individual SNPs, although, sometimes causally linked to trait variation, are highly redundant and cannot be readily used to annotate functional elements in the genome. Haplotypes aggregate genomic information over broad regions and are a powerful tool to link genomic variation to trait variation. However, they are usually too broad to be used to annotate regions less than a few megabases in length. By combining the mapping power of haplotypes, the high resolution of SNPs, and the intermediate resolution of chromatin states, we can begin to build mechanistic hypotheses that link genetic variation to variation in gene expression and physiology. Understanding the role that genetic variation plays in modifying the chromatin state landscape will be critical in making these links. Through this survey we are providing a rigorous resource that explores the connection between genetic variation and epigenetic variation. Researchers in the wider community can query the epigenetic landscape of the nine DO/CC inbred founders to identify candidate regulatory regions in genes of interest and generate mechanistic hypotheses linking genetic variation to gene expression.

## Materials and Methods

### Ethics Statement

All animal procedures followed Association for Assessment and Accreditation of Laboratory Animal Care guidelines and were approved by Institutional Animal Care and Use Committee (The Jackson Laboratory, Protocol AUS #04008).

### Inbred Mice

Three female mice from each of nine inbred strains were used. Eight of these strains (129S1/SvImJ, A/J, C57BL/6J, CAST/EiJ, NOD/ShiLtJ, NZO/HlLtJ, PWK/PhJ, and WSB/EiJ) are the eight strains that served as founders of the Collaborative Cross/Diversity Outbred mice (Chesler *et al*., 2008). The ninth strain, DBA/2J, will facilitate the interpretation of existing and forthcoming genetic mapping data obtained from the BxD recombinant inbred strain panel. Samples were harvested from the mice at 12 weeks of age.

#### Liver perfusion

To purify hepatocytes from the liver cell population, the mouse livers were perfused with 87 CDU/mL Liberase collagenase with 0.02% CaCl2 in Leffert’s buffer to digest the liver into a single-cell suspension, and then isolated using centrifugation.

We aliquoted 5 × 10^6^ cells for each RNA-Seq and bisulfite sequencing, and the rest were cross-linked for ChIP assays. Both aliquots were spun down at 200 rpm for 5 min, and resuspended in *1200μL* RTL+BME (for RNA-Seq) or frozen as a cell pellet in liquid nitrogen (for bisulfite sequencing). In the sample for ChIP-Seq, protein complexes were cross-linked to DNA using 37% formaldehyde in methanol. All cell samples were stored at −80°C until used (See Supplemental Methods for more detail).

#### Hepatocyte histone binding and gene expression assays

Hepatocyte samples were used in the following assays:

1. RNA-seq to quantify mRNA and long non-coding RNA expression, with approximately 30 million reads per sample.
2. Reduced-representation bisulfate sequencing to identify methylation states of approximately two million CpG sites in the genome. The average read depth was 20-30x.
3. Chromatin immunoprecipitation and sequencing to assess binding of the following histone marks:

a. H3K4me3 to map active promoters
b. H3K4me1 to identify active and poised enhancers
c. H3K27me3 to identify closed chromatin
d. H3K27ac, to identify actively used enhancers
e. A negative control (input chromatin)

Samples were sequenced with ~ 40 million reads per sample.

The samples for RNA-Seq in RTL+BME buffer were sent to The Jackson Laboratory Gene Expression Service for RNA extraction and library synthesis.

#### Histone chromatin immunoprecipitation assays

After extraction, hepatocyte cells were lysed to release the nuclei, spun down, and resuspended in 130ul MNase buffer with 1mM PMSF (Sigma, #78830) and 1x protease inhibitor cocktail (Roche) to prevent histone protein degradation. The samples were then digested with 15U of micrococcal nuclease (MNase), which digests the exposed DNA, but leaves the nucleosome-bound DNA intact. We confirmed digestion of nucleosomes into 150bp fragments with agarose gel. The digestion reaction was stopped with EDTA and samples were used immediately in the ChIP assay. The ChIP assay was performed with Dynabead Protein G beads and histone antibodies (H3K4me3: Millipore #07-473, H3K4me1: Millipore #07-436, H3K27me3: Millipore #07-449, H4K27ac: abcam ab4729). After binding to antibodies, samples were washed to remove unbound chromatin and then eluted with high-salt buffer and proteinase K to digest protein away from DNA-protein complexes. The DNA was purified using the Qiagen PCR purification kit. Quantification was performed using the Qubit quantification system (See Supplemental Methods).

### Diversity Outbred mice

We used previously published data from a population of 478 diversity outbred (DO) mice (Svenson *et al*., 2012). DO mice (JAX:DO) are available from The Jackson Laboratory (Bar Harbor, ME) (stock number 009376). The DO population included males and females from DO generations four through 11. Mice were randomly assigned to either a chow diet (6% fat by weight, LabDiet 5K52, LabDiet, Scott Distributing, Hudson, NH), or a high-fat, high-sucrose (HF/HS) diet (45% fat, 40% carbohydrates, and 15% protein) (Envigo Teklad TD.08811, Envigo, Madison, WI). Mice were maintained on this diet for 26 weeks.

#### Genotyping

All DO mice were genotyped as described in Svenson *et al*. (2012) (Svenson *et al*., 2012) using the Mouse Universal Genotyping Array (MUGA) (7854 markers), and the MegaMUGA (77,642 markers) (GeneSeek, Lincoln, NE). All animal procedures were approved by the Animal Care and Use Committee at The Jackson Laboratory (Animal Use Summary # 06006).

Founder haplotypes were inferred from SNPs using a Hidden Markov Model as described in Gattie *et al*. (2014) (Gatti *et al*., 2014). The MUGA and MegaMUGA arrays were merged to create a final set of evenly spaced 64,000 interpolated markers.

#### Tissue collection and gene expression

At euthenasia, whole livers were collected and gene expression was measured using RNA-Seq as described perviously (Chick *et al*., 2016; Tyler *et al*., 2017). Briefly, hepatocyte RNA was isolated using the Trizol Plus RNA extraction kit (Life Technologies), and 100-bp single-end reads were generated on the Illumina HiSeq 2000.

### Data Processing

#### Sequence processing

The raw sequencing data from both RNA-Seq and ChIP-Seq were put through the quality control program FastQC (0.11.5), and duplicate sequences were removed before downstream analysis.

#### Transcript quantification

Transcript sequences were aligned to strain-specific pseudo-genomes (*Chick et al*., 2016), which integrate SNPs and indels from each strain based on the GRCm38 mouse genome build. The B6 samples were aligned directly to the reference mouse genome. The pseudogenomes were created using g2gtools (http://churchill-lab.github.io/g2gtools/#overview). We used EMASE (https://github.com/churchill-lab/emase) to quantify the gene expression counts and DESeq2 vst transformation (Love *et al*., 2014) to normalize the gene expression data. We filtered out transcripts with less than 1 CPM in two or more replicates.

#### ChlP-Seq quantification

We used MACS 1.4.2 (Zhang *et al*., 2008) to identify peaks in the ChIP-Seq sequencing data, with a significance threshold of *p* ≤ 10^-5^. In order to compare peaks across strains, we converted the MACS output peak coordinates to common B6 coordinates using g2g tools (https://churchilllab.github.io/g2gtools/).

### Quantifying DNA methylation

RRBS data were processed using a bismark-based pipeline modified from (Thompson *et al*., 2018). The pipeline uses Trim Galore! 0.6.3 (https://www.bioinformatics.babraham.ac.uk/projects/trim_galore/) for QC, followed by the trimRRBSdiversityAdaptCustomers.py script from NuGen for trimming the diversity adapters. This script is available at: https://github.com/nugentechnologies/NuMetRRBS

All samples had comparable quality levels and no outstanding flags. Total number of reads was 45-90 million, with an average read length of about 50 bp. Quality scores were mostly above 30 (including error bars), with the average above 38. Duplication level was reduced to < 2 for about 95% of the sequences.

High quality reads were aligned to a custom strain pseudogenomes, using bowtie2 as implemented in Bismark 0.22 (Krueger and Andrews, 2011). The pseudogenomes were created by incorporating strain-specific SNPs and indels into the reference genome using g2gtools (https://github.com/churchilllab/g2gtools), allowing a more precise characterization of methylation patterns. Bismark methylation extractor tool was then used for creating a bed file of estimated methylation proportions for each animal, which was then translated to the reference mouse genome (GRCm38) coordinates using g2gtools. Unlike other liftover tools, g2gtools does not throw away alignments that land on indel regions. B6 samples were aligned directly to the reference mouse genome (mm10).

### Analysis of histone modifications

#### Identification of chromatin states

We used ChromHMM (1.22) (Ernst and Kellis, 2017) to identify chromatin states, which are unique combinations of the four chromatin modifications, for example, one state could consist of high levels of both H3K4me3 and H3K4me1, and low levels of the other two modifications. We conducted all subsequent analyses at the level of the chromatin state.

To ensure we were analyzing the most biologically meaningful chromatin states, we calculated chromatin states for all numbers of states between four and 16, which is the maximum number of states possible with four binary chromatin modifications (2^*n*^). We aligned states across the models by assigning each to one of the sixteen possible binary states using an emissions probability of 0.3 as the threshold for presence/absence of the histone mark. This threshold was used for comparison purposes only, and produced the most stable state estimates between models. We then investigated the stability of three features across all states: the emissions probabilities (Supp Fig1), the abundance of each state across transcribed genes (Supp Fig2), and the effect of each state on transcription (Supp Fig3). Methods for each of these analyses are described separately below. All measures were remarkably consistent across all models, but the 14-state model was characterized by a wide range of relatively abundant states with relatively strong effects on expression. We used this model for all subsequent analyses. For more details on how the different models were compared, see Supplemental Methods.

#### Genome distribution of chromatin states

We investigated genomic distributions of chromatin states in two ways. First, we used the ChromHMM function OverlapEnrichment to calculate enrichment of each state around known functional elements in the mouse genome. We analyzed the following features:

- Transcription start sites (TSS) - Annotations of TSS in the mouse genome were provided by RefSeq (O’Leary *et al*., 2016) and included with the release of ChromHMM, which we downloaded on December 9, 2019 (Ernst and Kellis, 2017).
- Transcription end sites (TES) - Annotations of TES in the mouse genome were provided by RefSeq and included with the release of ChromHMM.
- Transcription factor binding sites (TFBS) - We downloaded TFBS coordinates from OregAnno (Lesurf *et al*., 2016) using the UCSC genome browser (Kent *et al*., 2002) on May 4, 2021.
- Promoters - We downloaded promoter coordinates provided by the eukaryotic promoter database (Dreos *et al*., 2017,?), through the UCSC genome browser on April 26, 2021.
- Enhancers - We downloaded annotated enhancers provided by ChromHMM through the UCSC genome browser on April 26, 2021.
- Candidates of cis regulatory elements in the mouse genome (cCREs) - We downloaded cCRE annotations provided by ENCODE (Dunham *et al*., 2012; Moore *et al*., 2020) through the UCSC genome browser on April 26, 2021.
- CpG Islands - Annotations of CpG islands in the mouse genome were included with the release of ChromHMM.

In addition to these enrichments around individual elements, we also calculated chromatin state abundance relative to the main anatomical features of a gene. For each transcribed gene, we normalized the base pair positions to the length of the gene such that the transcription start site (TSS) was fixed at 0, and the transcription end site (TES) was fixed at 1. We also included 1000 bp upstream of the TSS and 1000 bp downstream of the TES, which were converted to values below 0 and above 1 respectively. To map chromatin states to the normalized positions, we binned the normalized positions into bins of relative position incremented by 0.1 and encompassing all upstream and downstream positions originally defined as 1kb up and downstream of the gene body. If a bin encompassed multiple positions in the gene, we assigned the mean value of the feature of interest to the bin. To avoid potential contamination from regulatory regions of nearby genes, we only included genes that were at least 2kb from their nearest neighbor, for a final set of 14,048 genes.

#### Chromatin state and gene expression

We calculated the effect of each chromatin state on gene expression. We did this both across genes and across strains. The across-gene analysis identified states that are associated with high expression and low expression within the hepatocytes and independent of strain. The across-strain analysis investigated whether variation in chromatin state across strains contributed to variation in gene expression across strains.

For each transcribed gene, we calculated the proportion of the gene body that was assigned to each chromatin state. We then fit a linear model separately for each state to calculate the effect of state proportion with gene expression:

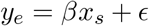

where *y_e_* is the rank normal scores (Conover, 1999) of the full transcriptome in a single inbred strain, and *x_s_* is the rank normal proportion of each gene that was assigned to state *s*. We fit this model for each strain and each state to yield one *β* coefficient with a 95% confidence interval. We fit the strains independently to better identify variation in chromatin state effects across strains. However, the effects were not different across strains (ANOVA *p* > 0.5), so we averaged the effects and confidence intervals across strains to yield one summary effect for each state. We further fit models for each state independently, rather than a multiple regression, because we were primarily interested in the marginal effects of each state for this study.

To calculate the effect of each chromatin state across strains, we first standardized transcript abundance across strains for each transcript. We also standardized the proportion of each chromatin state for each gene across strains. We then fit the same linear model, where *y_e_* was a rank normal vector concatenating all standardized expression levels across all strains, and *x_s_* was a rank normal vector concatenating all standardized state proportions across all strains. We fit the model for each state independently yielding a *β* coefficient and 95% confidence interval for each state.

In addition to calculating the effect of state proportion across the full gene body, we also performed the same calculations in a position-based manner. This second analysis yielded an effect of each state at multiple points along the gene body and a more nuanced view of the effect of each state.

### Analysis of DNA methylation

#### Creation of DNA methylome

We combined the DNA methylation data into a single methylome cataloging the methylated sites across all strains. For each site, we averaged the percent methylation across the three replicates in each strain. The final methylome contained 5,311,670 unique sites across the genome. Because methylated CpG sites can be fully methylated, unmethylated, or hemi-methylated, we rounded the average percent methylation at each site to the nearest 0, 50, or 100%.

#### Distribution of CpG sites

We used the enrichment function in ChromHMM described above to identify enrichment of CpG sites around functional elements in the mouse genome. We further performed a gene-based analysis of abundance similar to that in the chromatin states. As a function of relative position on the gene body, we calculated the density of CpG sites as the average distance to the next downstream CpG site, as well as the percent methylation at each site.

#### Effects of DNA methylation on gene expression

As with chromatin state, we assessed the effect of DNA methylation on gene expression both across genes and across strains. We used the same linear model described above, except that *y_s_* became the rank Z normalized percent methylation either across genes or across strains. Because the effect of DNA methylation on gene expression is well-known to be dependent on position, we only calculated a position-dependent effect on expression. We did not calculate the effect of percent methylation across the full gene on expression.

#### Imputation of genomic features in Diversity Outbred mice

To assess the extent to which chromatin state and DNA methylation explained local expression QTLs, we imputed local chromatin state and DNA methylation into the population of diversity outbred (DO) mice described above. We compared the effect of the imputed epigenetic features to imputed SNPs.

All imputations followed the same basic procedure: For each transcript, we identified the haplotype probabilities in the DO mice at the genetic marker nearest the gene transcription start site. This matrix held DO individuals in rows and DO founder haplotypes in columns (Supp. Fig. S4).

For each transcript, we also generated a three-dimensional array representing the genomic features (chromatin state, DNA methylation status, or SNP genotype) derived from the DO founders. This array held DO founders in rows, feature state in columns, and genomic position in the third dimension. The feature state for chromatin consisted of states one through 14, for SNPs feature state consisted of the genotypes A,C,G, and T.

We then multiplied the haplotype probabilities by each genomic feature array to obtain the imputed genomic feature for each DO mouse. This final array held DO individuals in rows, the genomic feature in the second dimension, and genomic position in the third dimension. This array is analagous to the genoprobs object in R/qtl2 (Broman *et al*., 2019). The genomic position dimension included all positions from 1 kb upstream of the TSS to 1 kb downstream of the TES. SNP data for the DO founders in mm10 coordinates were downloaded from the Sanger SNP database (Keane *et al*., 2011) on July 6, 2021.

To calculate the effect of each imputed genomic feature on gene expression in the DO population, we fit a linear model. From this linear model, we calculated the variance explained (*R*^2^) by each genomic feature, thereby relating gene expression in the DO to each position of the imputed feature in and around the gene body.

## Data Access

All raw and processed sequencing data generated in this study have been submitted to the NCBI Gene Expression Omnibus (GEO; https://www.ncbi.nlm.nih.gov/geo/) under accession number GSE213968.

Code to run the analyses in this study are at https://github.com/annaLtyler/Epigenetics_Manuscript.

## Acknowledgements

This work was funded by The Jackson Laboratory Director’s Innovation Fund and the National Institutes of Health grants R01 GM115518 (to G.W.C), GM070683 (to G.A.C), R35 GM133724 (to C.L.B.), and P30 CA034196. J.J.T. is a Scholar of the Leukemia & Lymphoma Society.

We gratefully acknowledge expert assistance from Genome Technologies, Gene Expression Services, and Information Technologies at The Jackson Laboratory.

## Supplemental Figure Legends

Figure S1: Comparison of emissions probabilities across all ChromHMM models. Each row contains data for a single ChromHMM model fit to the number of states indicated on either side of the row. Each set of four columns shows data for each of the four histone modifications. Each set is separated from the next by a column of gray for ease of visualization. The bottom row, the reference row, shows the ideal state that all model states are being compared to. Blue indicates absence of the histone mark and red indicates presence. For each ChromHMM model, each state was assigned to one of the reference states using an emissions probability of 0.3 as a threshold for presence of the histone modification. If a state was not present in the given model, the corresponding area is shown in gray. Emissions probabilities near 0 are shown in blue, and probabilities near 1 are shown in red. Orange and yellow indicate intermediate probabilities. Aligning the states across all models shows a remarkable stability in the emissions across models, seen as vertical bars of consistent color.

Figure S2: Comparison of state abundance across all ChromHMM models. The left-most column shows the annotation for each state. Unannotated states are marked with a dash. The binary heatmap indicates which histone modifications were present in each state: 1 indicates presence, and 0 indicates absence. The histone modifications are labeled at the bottom of each column. The continuous heatmap shows the abundance of each state (in rows) in each ChromHMM model (in columns). The abundance is the proportion of transcribed genes with the state present. Less abundant states are shaded blue, and more abundant states are shaded yellow, orange, and red. The number of states in the model is indicated at the bottom of each column. The black box highlights the model used in this study – the 14-state model. State abundance was remarkably stable across the different models.

Figure S3: Comparison of state effect across all ChromHMM models. This figure is identical to Figure S2, except that the cells in the continuous heatmap show the effect of each state on gene expression across all ChromHMM models. The effect was the *β* coefficient derived from a linear model. Similar to state abundance, the effects were remarkably stable across models.

Figure S4: Schematic for imputation of histone modifications into the DO mice. For a single transcript imputation was made by multiplying a three-dimensional array, containing chromatin state by strain by position, by a two-dimensional array, contatining haplotype probabilities by DO individual, to create a three-dimensional array, containing individual by position by chromatin state probability.

Figure S5: Effects of percent DNA methylation for all (A) CpG sites, (B) bimodal CpG sites, and (C) trimodal CpG sites. The *y*-axis in each panel shows the effect of variation in DNA methylation on gene expression across hepatocyte genes. The gray polygon shows the 95% confidence interval of the effect. Although there is a weak negative effect on transcription of DNA methylation across all sites, there was no effect when looking at trimodal sites alone.

